# Integral membrane protein, *anchor*, is expressed in the *Drosophila* insulin-producing cells and is a novel modulator of homeostatic behaviors, including sleep, feeding, and sedation

**DOI:** 10.64898/2026.04.22.720032

**Authors:** Emmanuelle Palmieri, Mia Coon, Adekemi Sobukunola, Laurie Sutton, Fernando J. Vonhoff

## Abstract

Integral membrane proteins (IMPs) are central regulators of cellular signaling and represent a major class of therapeutic targets. GPR155 (also known as LYCHOS), an evolutionarily conserved protein containing both transporter-like and GPCR-like domains, has recently emerged as a lysosomal nutrient sensor implicated in mTORC1 signaling. Despite its enriched expression in brain regions associated with reward processing, the *in vivo* neuronal and behavioral functions of GPR155 remain undefined. Here, we leverage the genetic tractability of *Drosophila melanogaster* to characterize the role of the GPR155 ortholog, *anchor*, in neural circuit function and behavior. Here, we demonstrate that pan-neuronal downregulation of *anchor* leads to significant alterations in multiple behaviors, including reduced feeding, disrupted light-dependent rhythmicity, decreased sleep, increased waking locomotor activity, and diminished sedation sensitivity to ethanol. We also selectively manipulated *anchor* expression in the neuroendocrine insulin-producing cells (IPCs), which phenocopied impaired rhythmicity and decreased ethanol sedation sensitivity observed in pan-neuronal manipulations, indicating that *anchor* function within IPCs is sufficient to modulate discrete behavioral outputs. Our results suggest that *anchor* regulates behavior in a sexually dimorphic manner as changes in ethanol sedation sensitivity were more penetrant in females, whereas altered feeding and ethanol preference was observed only in males. These findings establish a previously unrecognized role for *anchor* in the regulation of neuroendocrine signaling and behavior. Given the conservation of mTORC1 signaling and neuropeptidergic systems across species, this work provides mechanistic insight into how multifunctional IMPs integrate metabolic and environmental cues to influence complex behaviors, with potential implications for understanding the molecular basis of feeding, sleep regulation, and substance use disorders.

## Introduction

Nearly 25% of human genes encode integral membrane proteins (IMP)s^1^, proteins permanently embedded in cellular membranes, including G-Protein Coupled Receptors (GPCRs), ion channels, transporters, and cell adhesion molecules. This class of proteins plays important roles in cellular communication and integration of environmental information, which makes them immensely important drug targets. In fact, they are acted on by >50% of approved drugs by the U.S. Food and Drug Administration (FDA)^2^. Furthermore, different IMPs often interact, engaging in cross-oligomerization to adjust signaling and cellular outcomes. However, a unique one, GPR155, combines multiple types of IMPs into one. GPR155 is comprised of a physically-linked transporter-like region and a GPCR-like region^3–5^.

Recently, GPR155 has garnered attention not only for its structure, but also for its involvement in a major cell growth pathway, mechanistic Target of Rapamycin Complex 1 (mTORC1) signaling^6,7^. Specifically, *in vitro* findings indicate that GPR155 localizes to lysosomes where it integrates nutrient information in order to modulate mTORC1 signaling, leading researchers to give it an additional name, LYsosomal CHOlesterol Sensing protein (LYCHOS)^7^. While mTORC1 is ubiquitously expressed, GPR155 has a specific expression pattern, especially in the brain. In particular, *in situ* hybridization experiments have shown highest expression of GPR155 in the pyramidal cell layer of the hippocampus and in the dorsal striatum^8,9^. Given the well-studied association of these brain regions with reward learning as well as mTORC1’s role in encoding rewarding information, these findings suggest that GPR155 could play a role in numerous behavioral outputs, particularly reward-related behaviors.

GPR155 is well-conserved among vertebrates and invertebrates. Previous studies in *Drosophila melanogaster* have identified the *anchor* gene as the fly ortholog of GPR155^10^. *anchor* regulates wing development and gut homeostasis by interacting with Bone Morphogenic Pathway (BMP)^10^ and c-Jun N-terminal kinase (JNK)^11^ signaling, respectively. However, the role of *anchor* in behavior and in the nervous system remains unknown, although its expression in neurons has been confirmed by RNAseq^12,13^. Here, we further characterize a specific neuronal population expressing *anchor*, the insulin-producing cells (IPCs).

The IPCs are a cluster of 14 neurosecretory cells, analogous to mammalian pancreatic β-cells, and they produce *Drosophila* Insulin-Like Peptides (DILPs) 2, 3, and 5^14^. They also express drosulfakinins (DSKs), which are cholecystokinin-like peptides involved in satiation^15^. IPCs are located in the *pars intercerebralis* (PI), the neuroendocrine center of the fly brain and functional homolog of the mammalian hypothalamus. From there, IPC axons terminate in several regions, including the *corpora cardiaca* (CC), a gland essential for glucose regulation. The CC is functionally similar to the pituitary gland in vertebrates, and it contains endocrine cells similar to vertebrate pancreatic α-cells^16,17^. IPCs additionally project into the fly crop (a stomach-like organ), the foregut, the anterior midgut, and to the anterior aorta, from which DILPs enter the hemolymph (the fly circulatory fluid)^18–20^. Besides ethanol sensitivity^21^, *Drosophila* IPCs have been described to regulate numerous aspects of behavior, ranging from appetite and feeding to sleep and circadian rhythm^22,23^. Thus, we sought to investigate whether manipulation of *anchor* would affect behaviors associated with IPC activity.

Here, we present the first study demonstrating the neuronal role of the GPR155 ortholog, *anchor*, in behavioral regulation. We used *Drosophila* as a genetic model system to study the *in vivo* role of *anchor* in neuronal function and behavioral regulation and found that pan-neuronal downregulation of *anchor* reduces feeding, light-cue dependent rhythmicity, and sleep. Pan-neuronal downregulation of *anchor* also increases waking locomotor activity and decreases sensitivity to ethanol. Consistently, *anchor* knockdown, exclusively in IPCs, phenocopies several behavioral defects observed following pan-neuronal *anchor*-RNAi manipulations, including decreased ethanol sensitivity and rhythmicity. Together, our findings show that IPCs regulate specific behaviors through *anchor.* Our work also reveals functions of *anchor* that are independent of the IPCs, such as regulating feeding, locomotor activity and sleep. Overall, given high behavioral and mechanistic conservation of fundamental signaling pathways, such as mTORC1, insights gained here advance the understanding of molecular underpinnings of alcohol-related behavior, sleep, and feeding.

## Materials and Methods

### Fly stocks

Flies were kept at 25 °C in a 12/12 h light/dark incubator on standard plastic vials, using a regular cornmeal/molasses food as described before^24,25^. The following fly lines were obtained from the Bloomington *Drosophila* Stock Center: elav^C155^–GAL4; UAS-Dcr (RRID:BDSC_25750); UAS-*anchor*RNAi1 (RRID:BDSC_51463); Dilp2-GAL4 (RRID:BDSC_37516); *anchor*-GAL4 (RRID:BDSC_66861; created via enhancer trapping using a GAL4-containing P-element^26^); w; UAS-mCherry.nls (RRID:BDSC_38425); *w*^1118^ (RRID:BDSC_6326). The following fly lines were obtained from the Vienna *Drosophila* Resource Center: UAS-*anchor*-RNAi2 (VDRC #8532).

“PN RNAi1” refers to pan-neuronal *anchor* knockdown flies (using the elav^C155^;UAS-Dcr driver). Notably, PN-RNAi2 was not used in behavioral experiments to do a high rate of lethality with the majority of progeny unable to successfully eclose. “IPC RNAi1/2” refers to *anchor* knockdown in the IPCs (using the Dilp2-GAL4 driver), generated with either *anchor* RNAi1 or *anchor* RNAi2.

To generate UAS-RNAi1 and UAS-RNAi2 controls, RNAi1 and RNAi2 stocks, respectively, were crossed to *w*^1118^. To generate the PN GAL4 and the IPC GAL4 controls, the pan-neuronal driver and the IPC driver, respectively, were likewise crossed to *w*^1118^. *anchor* RNAi knockdowns were confirmed in our qPCR analysis as described below. RNAi manipulations were used in all behavioral assays. Flies used in experiments were separated by sex under brief CO_2_ exposure no less than 24 hours prior to undergoing behavioral testing, dissection, or head collection.

### Immunohistochemistry

To visualize *anchor*-expressing cells in the fly brain, brains from flies expressing UAS nuclear mCherry under the control of *anchor*-GAL4 were dissected and used for immunolabeling. For all immunohistochemistry (IHC) experiments, whole-brain dissections were taken from 1–10-day old male and female adult cross progeny. Dissections were performed in ice-cold phosphate buffered saline (PBS) and fixed in 4% paraformaldehyde, as previously described^27,28^. Brains were rinsed 3x with cold PBST for 10 min at room temperature, and tissue samples were blocked in 5% normal goat serum (NGS) for an hour. Then they were incubated for 24 hours in primary antibody at 4°C (1:1300 rabbit anti-Dilp2 gifted by Jan Veenstra (the University of Bordeaux); RRID:AB_2569969^29^) diluted in 0.5% PBST at 4°C on a rotator. The following day, the brains were rinsed 3x in cold PBST for 10 min at room temperature and then incubated in secondary antibody (1:400 Alexa Fluor 488-conjugated goat anti-rabbit IgG; Invitrogen, USA; Cat. No. A-11008) for 2 hours at room temperature. The brains were rinsed 3x in cold PBST for 10 min at room temperature, immediately mounted in glycerol, and imaged at a Zeiss LSM 900 confocal microscope, using a 40X oil immersion objective. Representative images are select Z-stack projection slices, processed using ImageJ Fiji.

### Ethanol sedation

Sensitivity to ethanol-induced sedation was assessed in <9-day old flies, using a vapor-chamber assay as previously described^24^ with minor modifications. Briefly, 10 flies were tested per chamber. Chambers were gently tapped every 5 minutes, and the number of sedated flies were counted. Flies were considered sedated when they experienced the loss-of-righting reflex (LORR). Sedation time 50 (ST50) values represent the time point at which 50% of flies are sedated, showing LORR^24^.

### Ethanol preference

Ethanol preference was assessed, using the FlyPAD^30^, a capacitor assay, consisting of an arena that measures fly “sips” at two different sites. Over a 1-hour period, animals were given a choice between sucrose solution at one site and sucrose solution supplemented with ethanol resulting in a final ethanol concentration of 10% at the other site. Final concentration of sucrose solutions at both sites was 5%. Food solutions contain 1% agarose, so they do not move significantly from their placement site. Flies were tested at 7-11 days old. The experiment was run within 1-3 hours of the beginning of the light cycle, and subjects were starved for an hour prior to the experiment. Three flies were tested at a time in each arena. Runs with 0 sips were excluded from data analysis to eliminate potential technical failure and unhealthy flies.

### Feeding

Consumption of sucrose solution was measured over 24 hours, using the CApillary FEeding (CAFE)^31^ assay with modifications. Flies were given access to a capillary tube containing 10% sucrose solution. Green dye was added to the solution to allow for measurement of the volume in the capillary tube, using a digital micrometer. The initial volume and the volume after the 24-hour feeding period were measured. Measurements were also taken from apparatuses containing no flies in order to calculate average volume lost from evaporation. This volume was subtracted from all runs of a given testing day in order to calculate the volume eaten. Each run consisted of a vial containing 5 flies and a suspended capillary tube full of the sucrose solution. The vial had holes in the bottom to allow for air flow and a mesh to prevent subject escape. Vials were suspended over but not touching water in order to raise humidity and prevent the sucrose solution from drying out. All experiments began within 1-3 hours into the light cycle, and all experiments took place in the temperature and light-controlled housing incubator, set to 25°C.

### Circadian rhythm and sleep

Sleep and rhythmicity measures were assessed in <10-day old male flies, using the TriKinetics *Drosophila* Activity Monitoring System (DAM; TriKinetics, Waltham, MA, USA) as we previously described^32^. The DAM recorded locomotor activity counts in 1 min bins for 4 days in 12-h light/12-h dark (LD) followed by 3-6 days in constant darkness (DD). In LD conditions, Zeitgeber time (ZT) is used to indicate the time in 24 h cycle, with ZT0 being the “lights on” time, ZT12 the “lights off” time. Subjective day and subjective night take place during the DD when there is no light cue. Respectively, they refer to the time in DD that corresponds to the LD 12-h light phase and the 12-h dark phase^33^.

Sleep and rhythmicity data collected with the DAM were processed using Visualization and ANalysis of timE SerieS dAta—*Drosophila* Activity Monitors (VANESSA-DAM, v1.0.3)^34^. Specifically, VANESSA-DAM-CRA was used for processing rhythmicity measures, including the rhythmicity power and period length, and generating actograms. Actograms were plotted for each day, while all other measures were averaged over the entire LD or entire DD experiment. The power–significance threshold value, often referred to as the relative power, indicates how much stronger the observed rhythmic signal is compared to the background noise level, and it is used as an indicator of rhythm robustness. Using the VANESSA-DAM software, under chi-square analysis of periodograms, for a given p-value (p < 0.05), we calculated a power threshold based on number of data points, variance, and distribution of power across the tested periods. The robustness of the rhythmicity was determined by computing the difference between the peak chi-squared value and the value expected by chance.

VANESSA-DAM-SA was used for processing sleep measures, including sleep fraction, nighttime sleep, daytime sleep, sleep fraction, and activity index. Sleep was defined as five or more consecutive minutes of inactivity^35–37^. Daytime sleep was classified as sleep occurring during the lights-on phase (Zeitgeber Time (ZT) 0–ZT12), while nighttime sleep was defined as sleep occurring during the lights-off phase (ZT12–ZT0). In order to examine possible locomotion defects, we calculated the activity index (or waking activity index), which is a measure of activity counts per minute. During DD Circadian Time (CT) is used to describe time because there is no light cue.

### Phototaxis assay

Light perception was assessed through a phototaxis assay in which flies cross from a dark area towards a light source. The apparatus consisted of a tube with a dark side and a light side separated by a door. The dark side was wrapped in aluminum foil, and flies were habituated there for at least 30 minutes prior to the start of the experiment. The light side was not wrapped in foil, and it was next to a lamp, providing a light cue. Dim red light was used for short periods of time for experimenter visibility. All other light sources in the experiment room were turned off or removed. At the start of the experiment, the door separating the dark side from the light side was opened, and flies were allowed for 5 seconds to cross into the light side. After 5 seconds, the door was closed, and the number of flies on the light side were counted. 5 flies were tested in each run.

### RT-qPCR

To verify that *anchor* transcripts were reduced and assess relative strength of knockdowns generated with RNAi1 and RNAi2, RT-qPCR was performed in male and female pan-neuronal *anchor* knockdown flies and corresponding controls as we previously described^24^. Whole-head samples from 5-10 day-old flies were used. Control and experimental flies were snap frozen in liquid nitrogen and heads were isolated. Total RNA was extracted using Quick-RNA MiniPrep (Zymo Research) according to manufacturer’s instructions and stored at −80°C. Total RNA was reverse transcribed using iScript cDNA Synthesis Kit (Bio-Rad) according to manufacturer’s instructions. Quantification of transcripts were assessed, using SYBR Green and the CFX Real-Time System C1000 Touch Thermal Cycler (Bio-Rad). *anchor* transcript levels were normalized to those of endogenous actin42A. Data were processed, using the *Δ Δ*C_q_ method and presented as fold change compared to PN GAL4 control flies. Experiments were performed in biological groups of 3-4, each of which was performed in technical triplicate. Each biological replicate consisted of 8 fly heads. Primer pair sequences targeting actin were previously described^38^. Sequences for actin primers were (1) forward: GCGTCGGTCAATTCAATCTT and (2) Reverse: AAGCTGCAACCTCTTCGTCA. Sequences for the primer mix targeting *anchor* were (1) forward: GGGGCGTATCCATGAACAACT and (2) reverse: TTGAAGCGTCCGGCAATGTAA. Primers were designed for *anchor*, NCBI Reference Sequence: NM_001275066.1, using NCBI Primer-BLAST.

### Statistical analysis and graphing

GraphPad Prism was used for all statistical analysis and graphing. Statistical significance was determined in all measures, using one-way ANOVA with multiple comparisons. In all graphs, statistical significance is shown by ****p < 0.0001, ***p < 0.001, **p < 0.01, and *p < 0.05.

## RESULTS

### All 14 IPCs express reporter under control of *anchor*-GAL4

In order to study the role of *anchor* in the fly brain, our first step was to confirm the pattern of its neuronal expression. The Fly Atlas Brain Project^12,13^ provides a publicly accessible RNAseq dataset, indicating that *anchor* is expressed in several glial and neuronal populations, including the insulin-producing cells (IPCs). Whole-brain dissections were performed in flies expressing nuclear mCherry under a GAL4 transgene inserted into the *anchor* locus that we used as a reporter of *anchor* expression. We observed *anchor* expression in several areas of the fly brain, particularly in a region located within the *pars intercerebralis* (PI) that contains neurosecretory cell clusters. One cell population within the PI is the IPCs, which are singly-identifiable, stereotypic cells with each fly having 14 of them^14^. Given that they are major neuroendocrine cells involved in the regulation of homeostatic behavior, we chose the IPCs as our cell population of interest in order to characterize expression of *anchor* in a specific cell population.

To confirm expression of *anchor* in the IPCs, dissected brains were immunostained with anti-*Drosophila* insulin-like peptide 2 (DILP2), an IPC marker^39^ (Figure 1A). We observed that all 14 DILP2-positive cells also expressed mCherry under the control of *anchor-*GAL4 (Figure 1B-F, Supplemental Figure 1). Notably, additional non-IPC nuclei were visibly labeled with mCherry, which are thought to belong to cells making up the nearby glial sheath, consistent with RNAseq data indicating *anchor* transcript expression in glial cells^12,13^.

**Figure 1:**
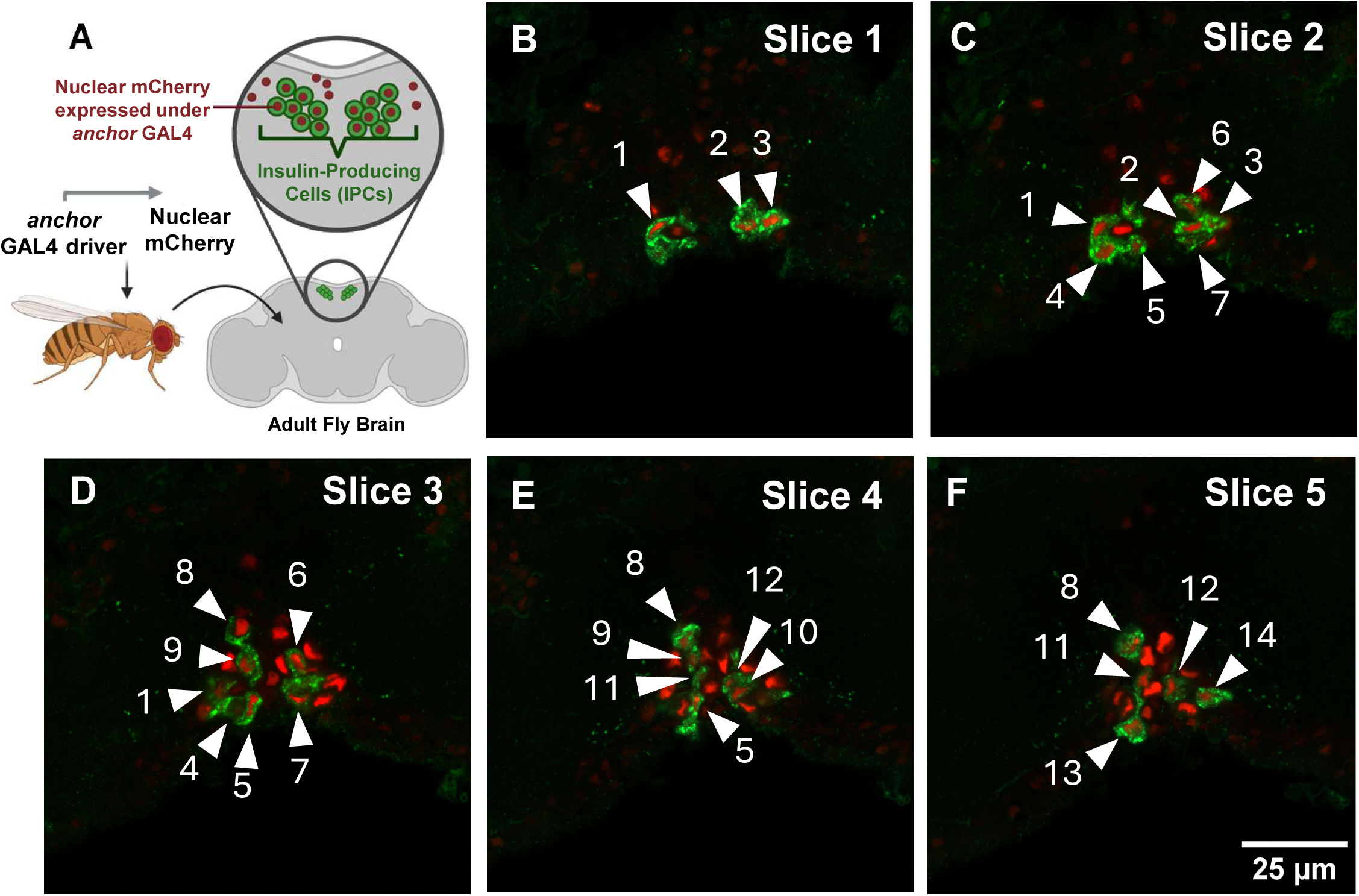
anchor reporter is expressed in all 14 of the insulin-producing cells (IPCs). A) Schematic of Immunohistochemistry experiment, using anti-DILP2 antibody (green) to image the insulin-producing cells (IPC)s in anchor GAL4 x nuclear mCherry (red) progeny. Made in Biorender. B-F) Representative confocal images from single fly z-stack (optical slices 1-5, n = 3). Merge showing expression of anchor-positive nuclei in DILP2-expressing neurons. Each of the 14 DILP2-expressing IPCs indicated by arrowhead and numbered from 1-14. Un-merged images available in Supplemental Figure 1.

As previous literature has linked the IPCs to ethanol-induced sedation^21^, feeding^24^, and regulation of circadian rhythm and sleep^22^, we performed several behavioral assays to further characterize the effects of *anchor* dysregulation in the nervous system and exclusively in the IPCs.

### *anchor* in the IPCs mediates ethanol-induced sedation

In order to characterize the role of *anchor* in the regulation of behavior, we used a pan-neuronal *anchor* knockdown model, generated using one of two different UAS *anchor* RNAi lines. We henceforth refer to pan-neuronal *anchor* knockdown flies as PN RNAi1 and PN RNAi2, respectively. Whereas a large number of progeny was obtained for PN RNAi1 flies, high lethality was observed in PN RNAi2 flies. Therefore, PN RNAi1 flies were used for both behavioral and molecular experiments, whereas the few PN RNAi2 escapers were used for molecular experiments. *anchor* knockdown was confirmed in both PN RNAi1 and PN RNAi2 using qRT-PCR (Supplemental Figure 2A-D). Expression of *anchor* in both PN RNAi and PN RNAi2 is reduced compared to genetic control groups. Notably, as whole-head samples were used in qPCR measures, intact expression of *anchor* in other non-neuronal head structures likely contribute to observed fold change.

We initially explored the role of *anchor* in ethanol-related behavior, given that IPCs regulate response to ethanol-induced sedation^21^. Using an ethanol vapor chamber assay to assess sedation sensitivity to ethanol, we observed that female PN RNAi1 flies sedated significantly slower than both control lines at all tested concentrations (65%, 73%, and 85%) (Figure 2A-C). Males sedated at the same rate as control animals at the lowest concentration, 65% (Figure 2D), and at 73% they sedated slower than only one control line (Figure 2E). However, male PN RNAi1 flies sedated significantly slower than both control groups at the highest concentration of ethanol, 85% (Figure 2F). Together, these findings suggest that knockdown of *anchor* in all neurons reduces sensitivity to the sedative effects of ethanol in both sexes, and the effect that *anchor* reduction has on ethanol sensitivity is dose dependent in males.

**Figure 2:**
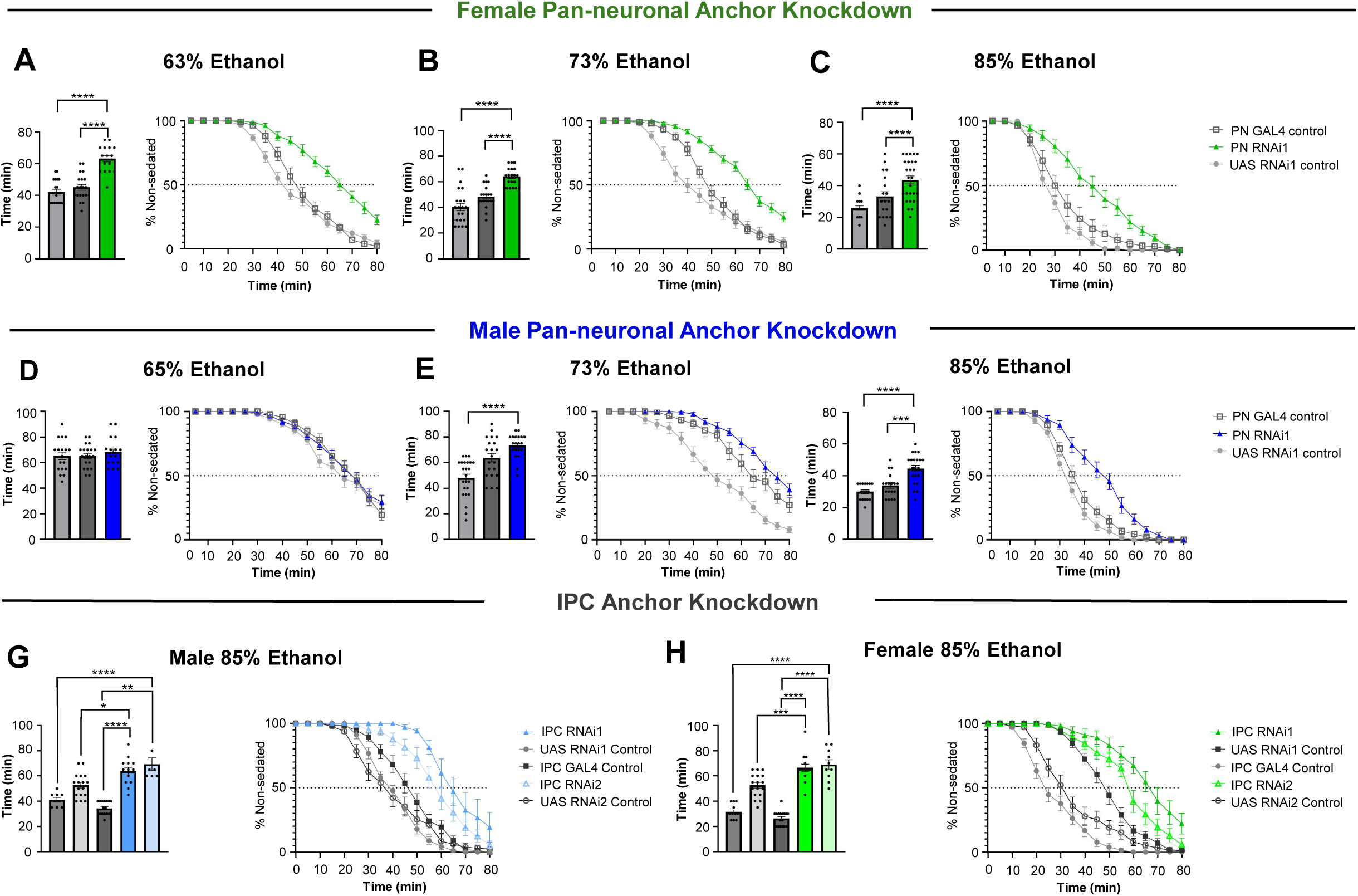
Pan-neuronal and IPC anchor knockdown increases time taken to sedate in the presence of ethanol vapor. A-C) Female and D-F) male pan-neuronal anchor knockdown and corresponding GAL4 and UAS control flies. 50% sedation point (ST50) with standard error of the mean (SEM) (left) and sedation over time (time course) (right) at 65%, 73%, and 85% ethanol concentration (N = 18-26). G) Male and H) female IPC anchor knockdown and corresponding GAL4 and UAS control flies. ST50 with SEM (left) and time course (right) at 85% ethanol concentration (N = 7-17). Results analyzed, using one-way ANOVA with multiple comparisons; *** p<0.001, **** p<0.0001. One run (N) includes 10 flies.

We next examined whether *anchor* function in a specific neuronal population was required for the regulation of ethanol sedation sensitivity. Given expression of *anchor* in the IPCs and evidence that the IPCs mediate ethanol-induced sedation, we hypothesized that knockdown of *anchor* in the IPCs would be sufficient to phenocopy the reduced ethanol sensitivity observed in the PN RNAi1 *anchor* knockdown flies. In order to test this hypothesis, the sedation assay was next performed using an IPC-specific driver and two different *anchor* RNAi lines (RNAi1 and RNAi2). Here, we refer to these knockdown lines as IPC-RNAi1 and IPC-RNAi2. Given the dose-dependent effect we observed in the PN RNAi1 males, the sedation assay was performed only at 85% ethanol. Consistently, in both males and females we observed that expression of either of the two *anchor* RNAi constructs in the IPCs induced reduced ethanol sensitivity compared to corresponding genetic controls (Figure 2G-H). These data suggests that *anchor* functions in the IPCs to regulate sedation sensitivity to ethanol exposure.

### *anchor* modulates ethanol preference and overall feeding

Because decreased sensitivity to ethanol is linked to increased ethanol preference^40^, we sought to determine whether *anchor* knockdown in the IPCs would also lead to higher preference for ethanol compared to corresponding genetic controls. To do so, we used the FlyPAD^30^, a capacitor system that tracks “sip” activity of the subject at two respective food sources presented at the same time (Figure 3A). We compared the number of sips at a 5% sucrose+10% ethanol solution to the number of sips at a control solution of 5% sucrose for 1 hour. We found that male IPC RNAi flies had significantly more sips at the solution supplemented with ethanol than the control solution, while corresponding control groups had a similar number of sips at the ethanol and the control solution (Figure 3B). This difference in preference was not observed in females (Figure 3E). For all groups, we observed no significant differences in the total number of sips taken and the sip duration during the 1-hour session in the FlyPAD (Figure 3C-D, F-G). These findings indicate that *anchor* knockdown in the IPCs is sufficient to induce increased ethanol preference in males.

**Figure 3:**
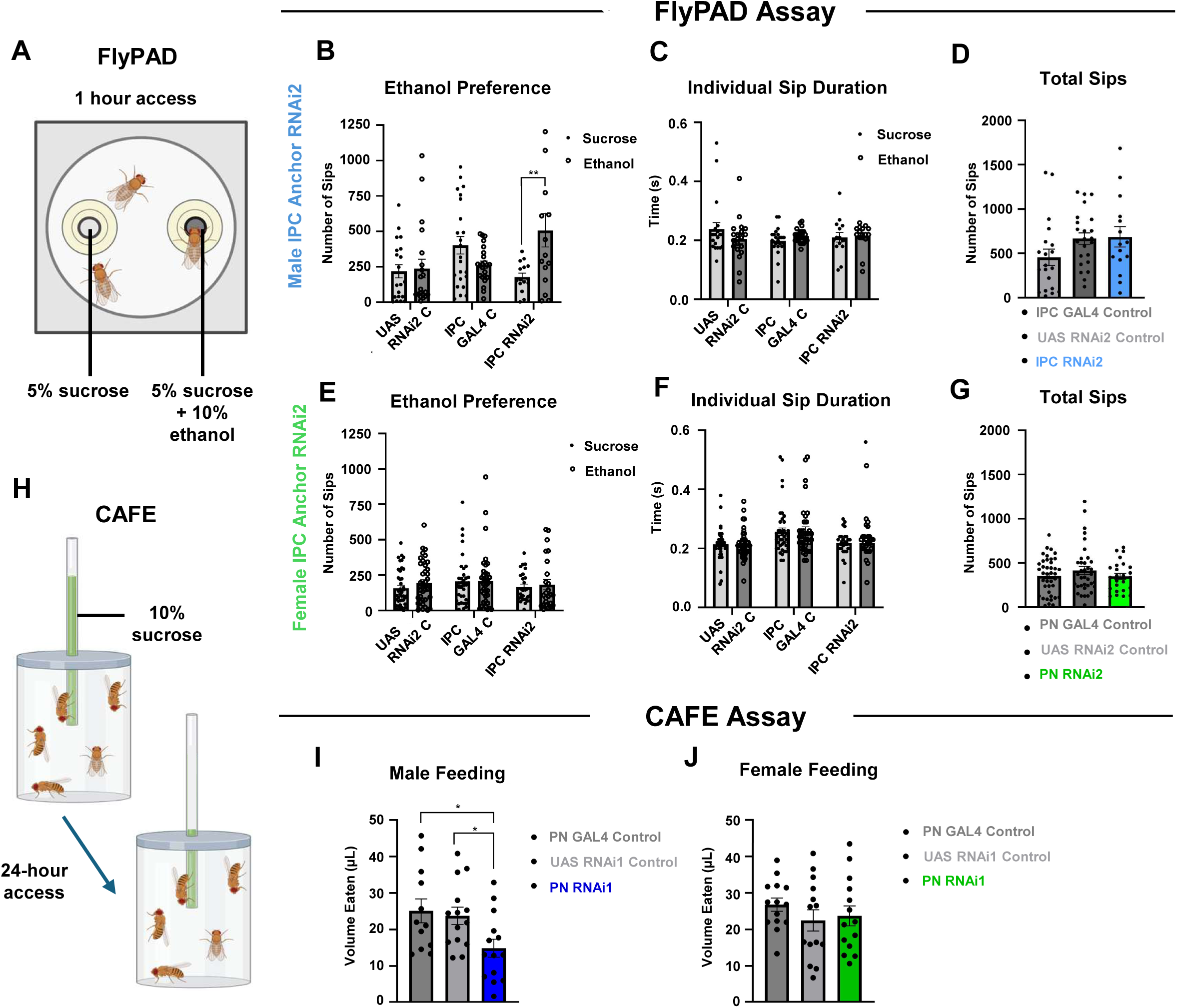
anchor knockdown increases ethanol preference and reduces overall food intake in males. A-G) Data for FlyPAD preference assay and H-J) for the CApillary FEeding (CAFE) assay. A) Schematic of the FlyPAD made in Biorender. B-D) Male (N = 15-23) and E-G) female (N = 25-40) IPC anchor RNAi2 and corresponding UAS and GAL4 control group total overall sips, individual sip duration, and ethanol preference. One run (N) includes 3 flies. H) Schematic of the CAFE assay made in Biorender. I) Male (N = 12-14) and J) female (N = 14) PN anchor RNAi1 and corresponding UAS and GAL4 control group volumes eaten over 24-hour period in the CAFE assay. One run (N) includes 5 flies. Results analyzed, using one-way ANOVA with multiple comparisons; ** p<0.01, **** p<0.0001. RNAi2C: UAS RNAi2 Control, IPC GC: IPC GAL4 Control

In order to rigorously assess the effect of *anchor* knockdown in feeding rate, we quantified baseline feeding over a longer period of time using the CApillary FEeding (CAFE)^31^ assay during 24-hour sessions. This method consists of a capillary tube filled with 10% sucrose solution (Figure 3H), which flies feed from during a 24-hour period, allowing us to quantify the volume consumed during an entire day. Data from the CAFE assay show that male PN RNAi1 flies consume a significantly lower overall volume of sucrose solution compared to corresponding genetic controls (Figure 3I). Female *anchor* knockdown flies, on the other hand, show no significant differences in total sucrose consumed (Figure 3J). These results indicate that when compared to control males, PN RNAi1 males feed less on sugar-containing foods but prefer ethanol-containing foods when given a choice. Overall, these results suggest that *anchor* may regulate feeding and ethanol preference in a sexually dimorphic manner.

### *anchor* modulates light-cue dependent rhythmicity, sleep, and waking locomotion, and the IPCs contribute to *anchor* modulation of rhythmicity

As the IPCs play a major role in regulating sleep/wake cycles^22^, we measured waking locomotion, sleep, and rhythmicity using the *Drosophila* Activity Monitor (DAM) ^32^. Specifically, the DAM records activity counts in minute bins, and we defined a sleep bout in our analyses as immobility for 5 or more minutes. Waking locomotion is calculated based on activity counts during non-sleep minutes. For 4 days flies were kept on their regular 12-hour light/12-hour dark (LD) cycle, followed by 3-6 days in constant darkness (DD), while their activity was recorded throughout these phases. The LD provides insight into regular behavior with light stimulus, while the DD gives information about the endogenous clock in the absence of light. First, a phototaxis assay was also performed in PN RNAi1 in order to determine whether *anchor* knockdown affects basic light cue-dependent responses that could confound observation of circadian and sleep behaviors (Supplemental Figure 2A). All groups responded normally to a light cue, indicating that *anchor* knockdown produces no detectable deficits in light detection and response (Supplemental Figure 2B).

During the LD, all flies showed the expected activity pattern, with a bout of high activity at the beginning of the lights-on cycle, followed by brief sleep bouts in the middle of the day, with a final bout of high activity around 8 hours into the light cycle, which ended when the lights turned off^32,37^. The PN RNAi1 exhibited higher waking activity, lower LD rhythmicity (period power), and reduced sleeping time during the light cycle than corresponding controls (Figure 4A-D, F). However, all groups (the PN RNAi1 and corresponding controls) slept a similar amount during the dark cycle of the LD and had similar period length (Figure 4E, G). These findings indicate that *anchor* knockdown alters the baseline locomotion, LD rhythmicity, and daytime sleep.

**Figure 4:**
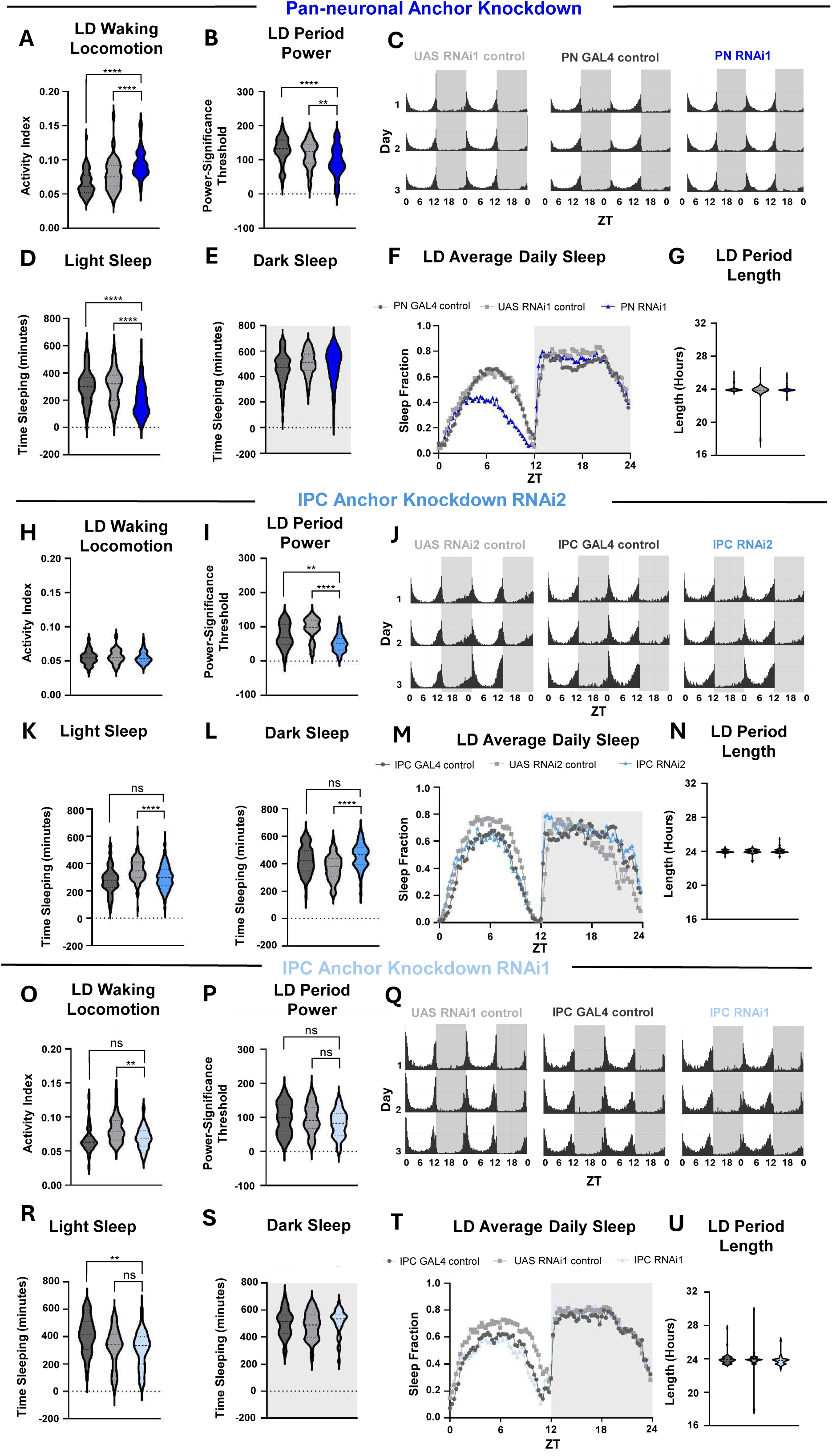
Pan-neuronal anchor knockdown increases waking activity, decreases daytime sleep, and dysregulates rhythmicity during LD, and IPCs contribute to anchor mediated rhythm dysregulation but not sleep or activity. A-G) Rhythmicity and sleep data for male pan-neuronal anchor knockdown and corresponding UAS and GAL4 control groups (n = 64-84). A) Activity during the LD. B) Period power given by the significance – threshold value. C) Actograms for each day of LD condition. D) Sleep during the daytime (12-h light). E) Sleep during the nighttime (12-h dark). F) Average daily fraction of time spent sleeping per hour throughout LD conditions. G) Average length of period. H-N) Rhythmicity and sleep data for IPC anchor knockdown RNAi2 and corresponding UAS and GAL4 control groups (n = 25-42). O-U) IPC anchor knockdown RNAi1 and corresponding UAS and GAL4 control groups (n = 31-50). Results analyzed using one-way ANOVA with multiple comparisons, ** p<0.01, **** p<0.0001.

To determine whether dysregulation of *anchor* specifically in the IPCs affects these parameters, experiments were conducted in the IPC-RNAi1 and IPC-RNAi2 and compared to their corresponding controls. Interestingly, we observed fully recapitulated deficits only in LD rhythmicity for IPC-RNAi2 (Figure 4I), whereas a partial effect on sleep and no significant differences in activity was observed between the IPC-RNAi2 and controls (Figure 4H, J-N). Notably, there were no fully recapitulated differences in any parameters for IPC-RNAi1 flies and corresponding controls (Figure 4O-U), which only show a partial effect on activity and sleep. The combination of decreased rhythmicity and normal activity levels suggests that *anchor* modulates LD rhythm independently of locomotion. Additionally, these findings suggest that, while the knockdown of *anchor* in the IPCs is sufficient to cause deficits in LD rhythmicity, it is not sufficient to reproduce the alterations in locomotion and in sleep observed in the PN-RNAi1. One explanation for this could be that knockdown of *anchor* in other, non-IPC cell populations, underlies the phenotypes observed in the PN-RNAi1. Together, these findings suggest that *anchor* functions in IPCs to regulate LD rhythmicity as well as in multiple cell populations to regulate locomotion and sleep.

### *anchor* mediation of rhythm, activity, and sleep is light-cue dependent

While behavior during the LD was used to understand *anchor’s* role in rhythm and sleep under normal daily conditions, we subsequently measured fly behavior under constant darkness to determine whether *anchor* knockdown causes observed phenotypes by affecting the endogenous clock or whether its role is light-dependent. The 4-day period of LD was followed, therefore, by 6 days of 24-hour darkness (DD).

Across all tested *anchor* knockdown groups and corresponding controls, we observe that none of the *anchor* RNAi-expressing groups have significant behavioral changes compared to both corresponding genetic controls. Especially notable is that none of the phenotypes affected by pan-neuronal RNAi1 expression during the LD (Figure 4) persist fully in the DD (Figure 5A-G). Sleep, rhythmicity, and activity are unaffected by *anchor* knockdown status during the DD. Likewise, in both IPC *anchor* knockdown groups there is no observable effect on any measured parameters (Figure 5H-U). Therefore, these findings indicate that the role *anchor* plays in rhythmicity, waking activity, and sleep is light dependent.

**Figure 5:**
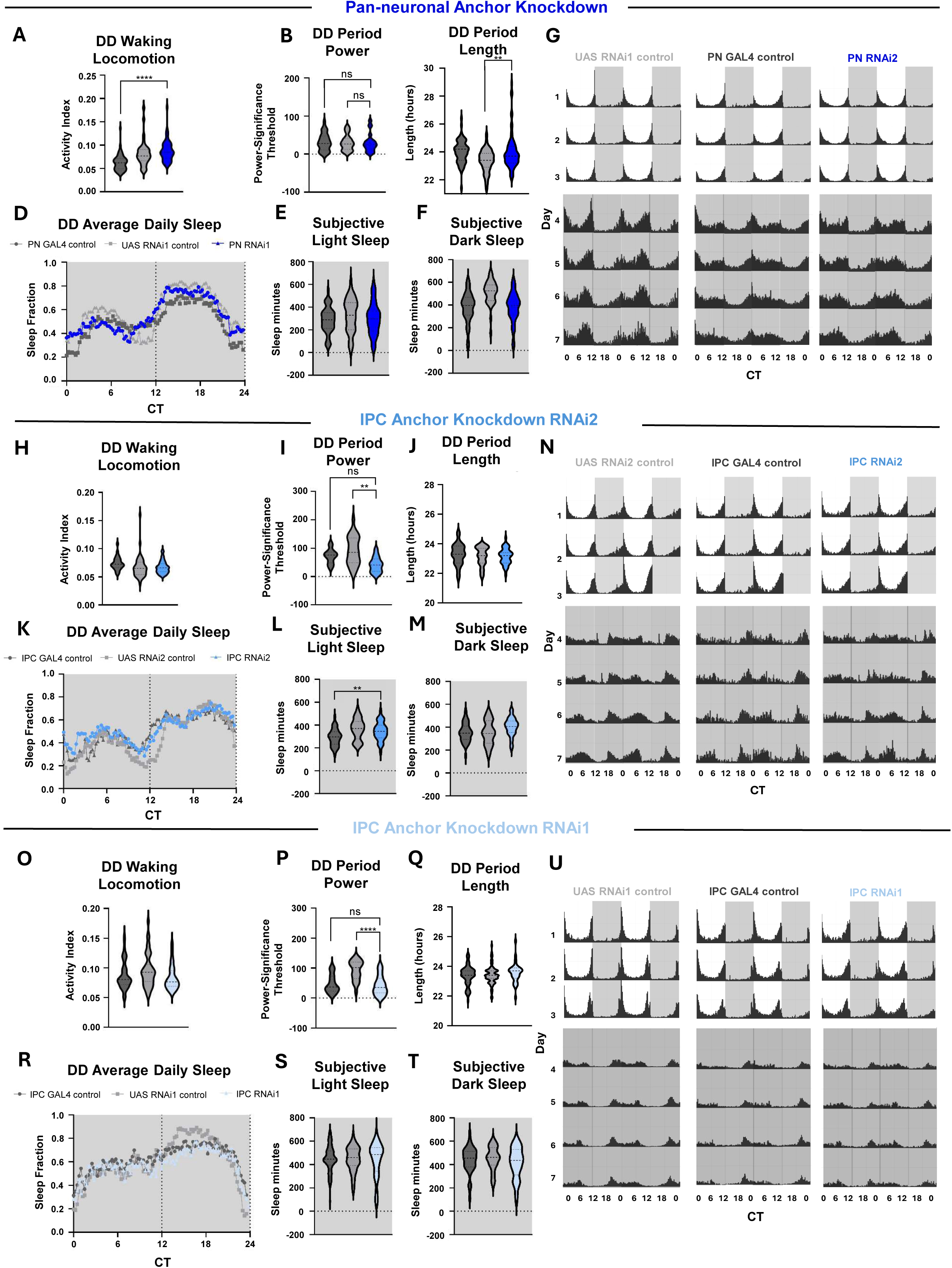
Under constant darkness, anchor knockdown produces no or only partial deficits in activity, sleep, and circadian rhythm. A-G) Rhythmicity and sleep data for male pan-neuronal anchor knockdown and corresponding UAS and GAL4 control groups (n = 30-51). A) Activity during the DD. B) Period power given by the significance – threshold value. C) Actograms for each day of DD condition. D) Sleep during subjective daytime (0h-12h CT). E) Sleep during subjective nighttime (12h-0h CT). F) Average daily fraction of time spent sleeping per hour throughout DD conditions. G) Average length of period. H-N) Rhythmicity and sleep data for IPC anchor knockdown RNAi2 and corresponding UAS and GAL4 control groups (n = 7-27). O-U) IPC anchor knockdown RNAi1 and corresponding UAS and GAL4 control groups (n = 14-22). Results analyzed using one-way ANOVA with multiple comparisons, ** p<0.01, **** p<0.0001.

## DISCUSSION

### First description of GPR155/anchor function in the nervous system indicates regulation of homeostatic behavior in a sexually dimorphic manner

While *anchor* in *Drosophila* has been previously associated with wing development and gut homeostasis^10,11^, to our knowledge this study represents the first description of the role of *anchor* in the nervous system. Our results indicate that *anchor* function in neurons is required to regulate homeostatic behaviors such as feeding, light cue-dependent rhythmicity, sleep, ethanol sedation sensitivity and ethanol preference. Interestingly, our results suggest that *anchor* is regulating behavior in a sexually dimorphic manner. The effect of pan-neuronal *anchor* downregulation on ethanol sedation sensitivity was more penetrant in females than in males. Whereas pan-neuronal *anchor* knockdown in females showed a significant decrease in sedation sensitivity at all tested concentrations (65%, 73% and 85%), PN RNAi1 males showed a significant effect only at 85%. This difference may be due to sex-specific expression levels of *anchor*, as RNAseq data indicates slightly higher expression levels of *anchor* in males than females, particularly in the brain^12,13^. Therefore, in contrast to males, the lower *anchor* levels in females may make them more susceptible to knockdown manipulations by reaching a low enough range able to cause observable phenotypes. Furthermore, previous publications have reported sexually dimorphic responses to ethanol exposure in *Drosophila*^41^, including our recent study showing juvenile hormone-dependent maturation of ethanol attraction^42^ and our sex-reversal study showing that feminization of *Corazonin* neurons in males led to female-like responses^24^. Whether *anchor* interacts with or is part of sex-specific neuronal networks or molecular pathways remains unknown.

Sex differences were additionally observed in both the feeding and the ethanol preference assay. Male *anchor* knockdown flies have increased ethanol preference and reduced sucrose consumption over a 24-hour period, while female *anchor* knockdown flies show neither of these differences. While we suggest that lower levels of *anchor* make females more susceptible to *anchor* knockdown in the sedation assay, here we propose that mating status affects baseline feeding behavior disproportionately to males. Females may have increased nutrient demands from egg-laying that mask potential differences between genetic control and *anchor* knockdown feeding and ethanol preference. Likewise, females were not tested in sleep and rhythmicity measures in this study. To solidify conclusions about the dimorphic nature of *anchor* effects on behavior as well as the role of *anchor* in interactions between mating status and behavior, virgin male and female subjects will be tested in measures of feeding and rhythmicity and sleep in future studies.

### Cellular characterization of GPR155/anchor expression and function

Our results provide experimental evidence that *anchor* is expressed in all of the IPCs in the brain. IPCs are a cluster of 14 neurosecretory cells, analogous to mammalian pancreatic β-cells, based on similar expression pattern of genes, and they are expressed in the PI of the fly brain, which is analogous to the mammalian hypothalamus. IPCs produce *Drosophila* Insulin-Like Peptides (DILPs) 2, 3, and 5^14^ in addition to expressing drosulfakinins (DSKs), which are cholecystokinin-like peptides^15^. In mammals, insulin is classically thought to be produced in β-cells of the pancreas, but mounting evidence reveals that insulin is also produced in several mammalian brain regions, including the hippocampus, cortex, and hypothalamus^43^. While GPR155 expression in pancreatic β-cells has not been reported, *in situ* experiments show that it is expressed in these brain regions. Mechanistically, GPR155 and *anchor* could have a similar behavioral role through action on insulin production via mTORC1 signaling (discussed below), though whether GPR155 has a similar behavioral role, whether *anchor* and GPR155 modulate mTORC1 signaling *in vivo,* and whether they act upstream of insulin remains to be studied.

Despite anatomical support for behavioral conservation of GPR155 and *anchor*, there are also differences in the expression patterns of *anchor* and GPR155. ^8,9^ For instance, GPR155 is highly expressed in the mammalian striatum. GPR155 is, in fact, a marker for a striatal subregion, the dorsolateral striatum (DLS)^8^. As the DLS is associated with habitual aspects of motor control^44^, this GPR155-expressing structure seems to regulate different behaviors from *Drosophila* IPCs.

Multiple studies have shown that IPCs regulate appetite and feeding. *In vivo* activation of IPCs in starved flies leads to satiety-like behavior^23^, and flies with depleted IPC DSKs consume significantly more food than controls, suggesting that DSKs suppress appetite acting as a hormonal satiety signal^15^. Based on the observed reduction in feeding in *anchor*-RNAi flies, our results suggest that *anchor* knockdown may phenocopy IPC activation and DSK release. Furthermore, IPCs also modulate sleep. Mutations in most DILPs decrease total sleep^45^, and knockdown of the Insulin Receptor (InR) and downstream molecules shorten circadian period and lower its amplitude^22^. However, expression of *anchor*-RNAi exclusively in IPCs did not affect sleep, but it affected light-dependent rhythmicity. Therefore, while *anchor* in the IPCs modulates LD rhythmicity, *anchor*-dependent changes in sleep and activity likely involve other neuronal populations, potentially photoreceptor neurons or ion transport peptide (ITP)-expressing neurons, which express *anchor*^46^. Future studies will examine the role of *anchor* in different neuronal populations besides the IPC-specific roles described here.

Lastly, impaired IPC function and reduced InR signaling in *Drosophila* increase sensitivity to ethanol^21^. By contrast, our results indicate that IPC-specific knockdown of *anchor* reduced sensitivity to the acute sedating effects of ethanol, consistent again with the idea that *anchor* knockdown may phenocopy IPC activation. As discussed below, whether *anchor* regulates the production or release of insulin remains to be answered. Lastly, since minimal anatomical conservation is observed in the distinct expression pattern of *anchor* and GPR155 in the fly and mouse brain, respectively, identification of molecular interactions downstream of *anchor*/GPR155 may reveal evolutionary conserved mechanisms (see below).

### Molecular mechanisms underlying GPR155/anchor-dependent regulation of behavior

Though not well studied in *Drosophila*, mTORC1 regulates insulin production in mammals through a negative feedback loop.^47^ *In vitro* mTORC1 signaling is reduced by GPR155 knockout because GPR155 cannot sequester the mTORC1 inhibitor, GATOR1 (guanosine triphosphatase-activating protein) via the lysosomal cholesterol signaling (LYCHOS) effector domain (LED) identified on GPR155^7^. In *Drosophila*, changes in mTOR signaling in IPCs are linked to altered production of insulin, which is linked to changes in feeding, ethanol sensitivity, and circadian rhythm. Therefore, a possible mechanism underlying the phenotypes observed in this study involves positioning *anchor* upstream of mTOR, which would in turn regulate insulin signaling. Future studies will assess whether *anchor* manipulations lead to changes in mTOR- and insulin-dependent signaling in a similar way as described in mammals.

Our results linking *anchor* function and ethanol-dependent behaviors in flies are consistent with mammalian studies showing that GPR155 is downregulated in the amygdala of rats that exhibit high ethanol preference^48^ as well as a established role of mTORC1 in addiction. Through its role in synaptic growth, striatal mTORC1 is thought to encode reinforcing properties of substances such as ethanol. In fact even during first-time exposure, ethanol triggers mTORC1-dependent plasticity in the striatum^49^. These findings underscore the need to better understand the link between GPR155/ *anchor*, mTORC1 signaling, and plasticity.

Moreover, the study of GPR155*/anchor* in synaptic plasticity and function may be relevant to understanding disease-associated phenotypes. Genome-Wide Association Studies (GWAS) have linked GPR155 dysregulation to neurological pathologies, including autism spectrum disorder^50^ and Huntington’s disease^51^. Altered GPR155 levels were observed in lymphoblastoid cells derived from human subjects with autism^50^ as well as in the caudate nucleus of the brains of subjects with Huntington’s disease^51^. Importantly, striatal GPR155 expression levels were found altered in a mouse model of Huntington’s disease as well as in a mouse model of Parkinson’s disease^52^. These findings suggest GPR155 could play an important role in synaptic function and in development of pathologies, but no studies have explored the direct manipulation of GPR155 *in vivo*. In particular, disrupted synaptic refinement has been associated with autism, as an increase in synaptic contacts is observed in brains of autistic patients^53^ as well as in *Drosophila* studies of autism-associated genes^32^. In *Drosophila*, mTOR signaling has been involved in synaptic plasticity^54^ but, although molecules associated with synaptic refinement such as CaMKI^32^, CaMKII^55,56^, and PKA^57^ are known to interact with mTOR^58,59^, a direct role of mTOR in synaptic refinement has not yet been confirmed^53^. Future research will test whether defects in synaptic refinement are observed following *anchor* manipulations via interactions with mTOR and the identified calcium- or cAMP-dependent refinement pathway^53^, elucidating GPR155/*anchor* possible role in plasticity and disease-associated pathways.

Besides mTOR, *anchor* has been previously described to interact with BMP^10^ and JNK^11^ signaling during wing development and gut homeostasis, respectively. Components of the BMP pathway have been not only associated with regulating feeding, sleep and circadian rhythms^60,61^, but also hypothesized to respond to ethanol exposure^62^. Similarly, JNK signaling has been involved in sleep regulation in flies^63^ as well as in regulating ethanol-dependent behaviors in flies, mice, and *C. elegans*^64,65^. As both BMP^66–69^ and JNK^70–72^ signaling are well-known regulators of synaptic plasticity, future research will determine whether behavioral phenotypes described here are a consequence of anatomical defects downstream of aberrant interactions between *anchor*, BMP and JNK signaling pathways.

## Acknowledgments

We would like to thank Jan Veenstra (University of Bordeaux) for kindly gifting us DILP2 antibody. We would also like to thank Arijit Ghosh for advice and guidance in our circadian and sleep analyses.

---

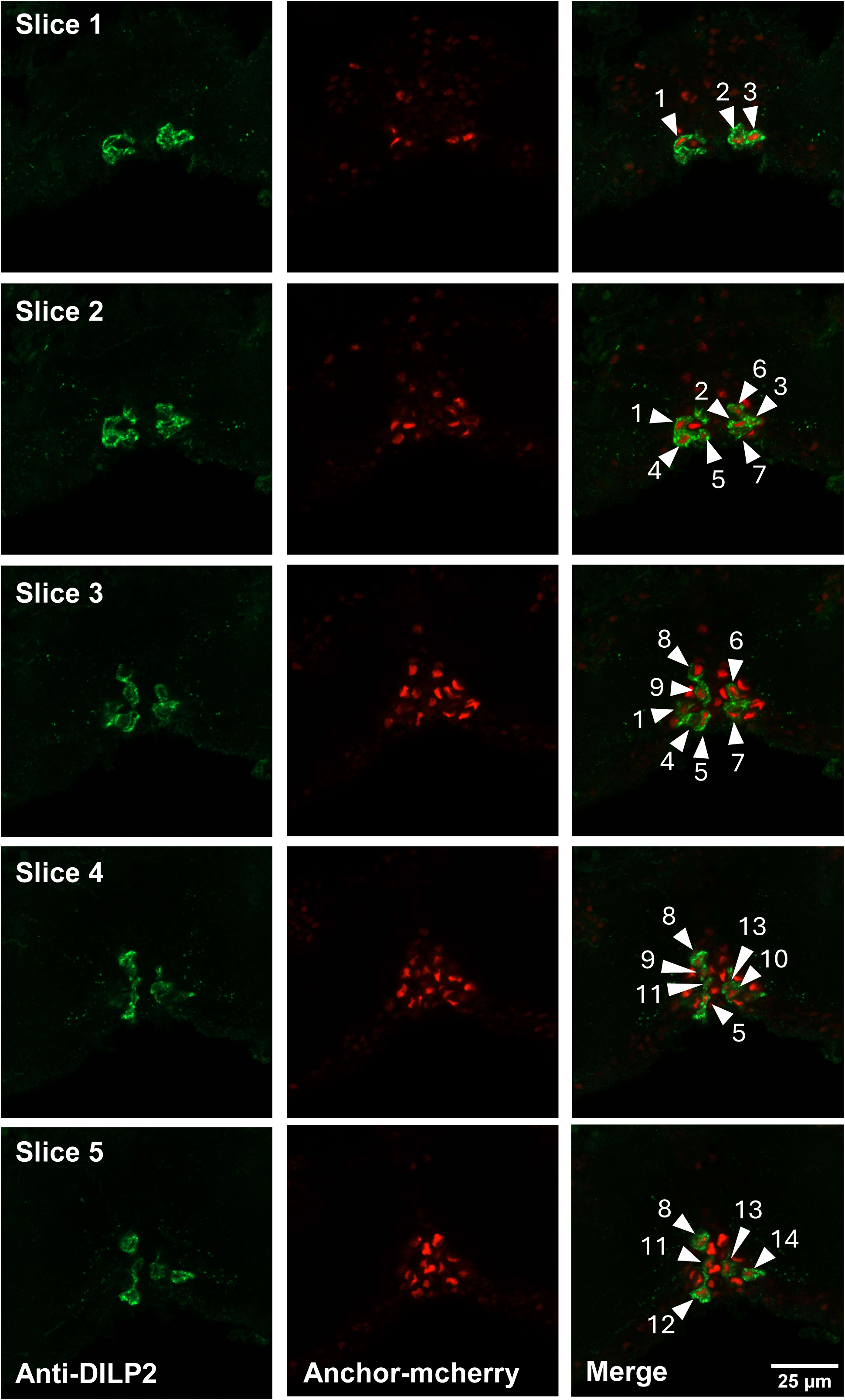

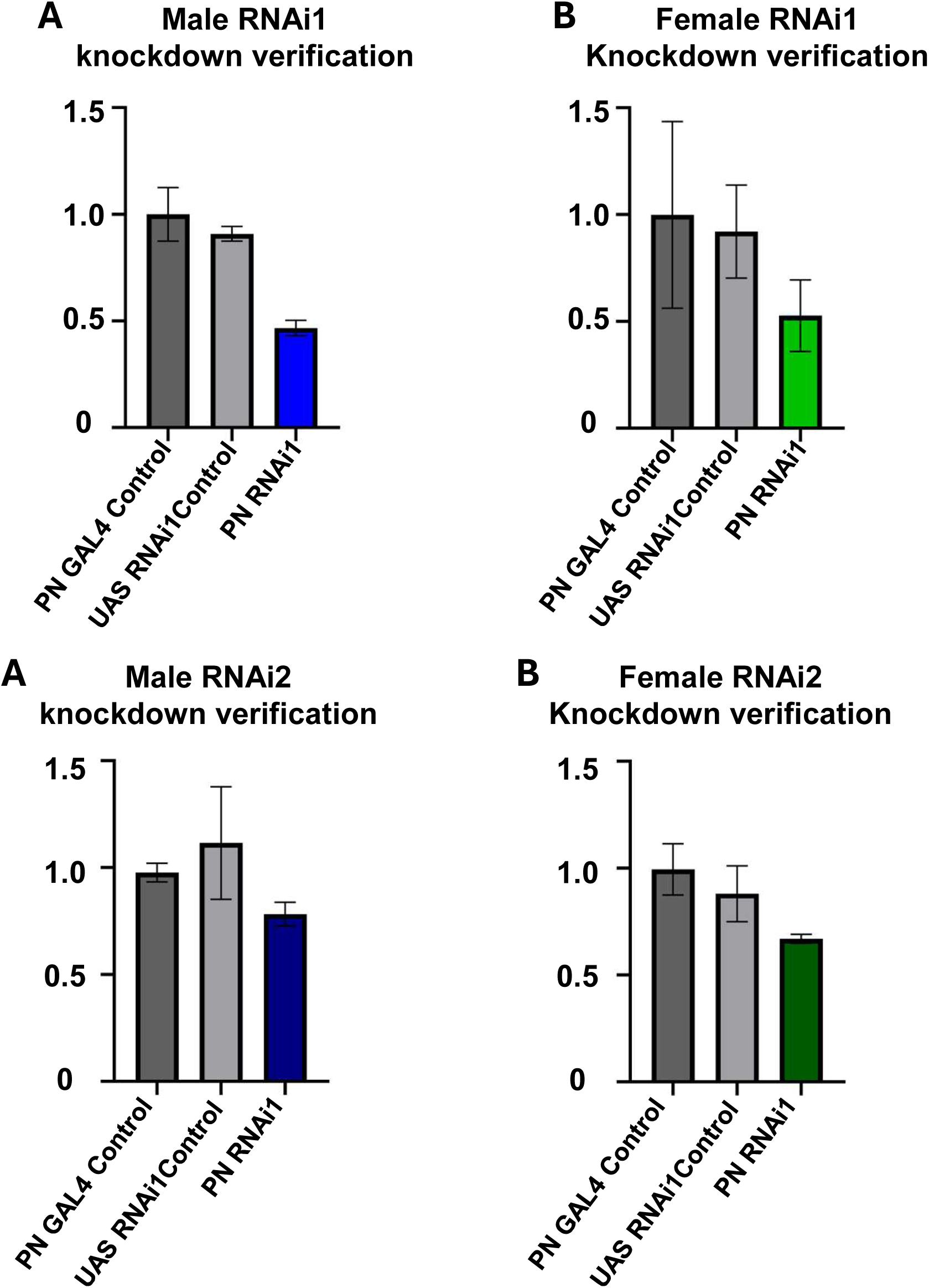

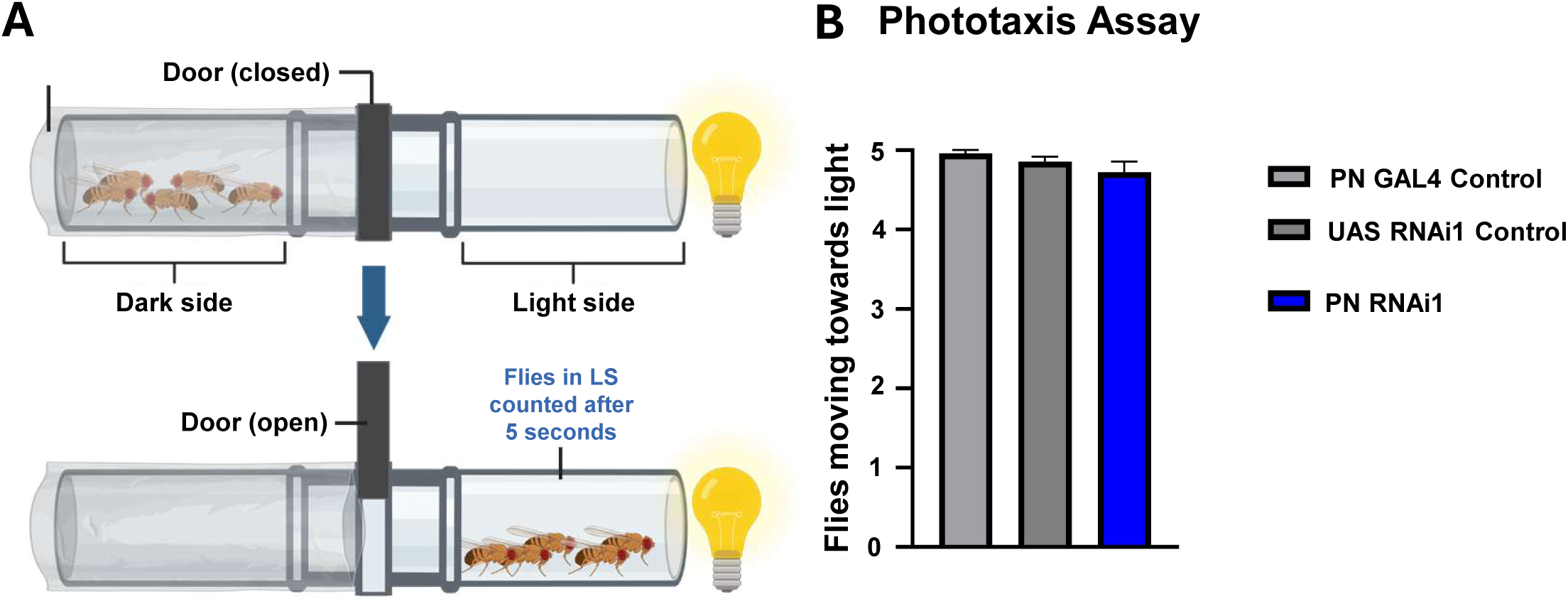

## REFERENCES

1. Fagerberg, L., Jonasson, K., Von Heijne, G., Uhlen, M. & Berglund, L. Prediction of the human membrane proteome. 1141–1149 (2010) 10.1002/pmic.200900258.

2. Yıldırım, M. A., Goh, K.-I., Cusick, M. E., Barabási, A.-L. & Vidal, M. Drug—target network. Nat Biotechnol 25, 1119–1126 (2007).

3. Sharma, M. et al. Cryo-EM structures of human GPR155 elucidate its regulatory and transport mechanisms. 2024.09.24.614577 Preprint at 10.1101/2024.09.24.614577 (2024).

4. Bayly-Jones, C. et al. LYCHOS is a human hybrid of a plant-like PIN transporter and a GPCR. Nature 634, 1238–1244 (2024).

5. Xiong, Q. et al. Molecular architecture of human LYCHOS involved in lysosomal cholesterol signaling. Nat Struct Mol Biol 32, 905–913 (2025).

6. Yu, S. et al. Structural insights into cholesterol sensing by the LYCHOS-mTORC1 pathway. Preprint at 10.1101/2024.11.21.624772 (2024).

7. Shin, H. R. et al. Lysosomal GPCR-like protein LYCHOS signals cholesterol sufficiency to mTORC1. Science 377, 1290–1298 (2022).

8. Märtin, A. et al. A Spatiomolecular Map of the Striatum. Cell Rep 29, 4320–4333.e5 (2019).

9. Trifonov, S. et al. GPR155: Gene organization, multiple mRNA splice variants and expression in mouse central nervous system. Biochemical and Biophysical Research Communications 398, 19–25 (2010).

10. Wang, X. C., Liu, Z. & Jin, L. H. Anchor negatively regulates BMP signalling to control *Drosophila* wing development. European Journal of Cell Biology 97, 308–317 (2018).

11. Wang, J., Liu, Q., Gong, Y. & Jin, L. H. Anchor maintains gut homeostasis by restricting the JNK and Notch pathways in Drosophila. Journal of Insect Physiology 134, 104309 (2021).

12. Leader, D. P., Krause, S. A., Pandit, A., Davies, S. A. & Dow, J. A. T. FlyAtlas 2: a new version of the Drosophila melanogaster expression atlas with RNA-Seq, miRNA-Seq and sex-specific data. Nucleic Acids Research 46, D809–D815 (2018).

13. Krause, S. A., Overend, G., Dow, J. A. T. & Leader, D. P. FlyAtlas 2 in 2022: enhancements to the *Drosophila melanogaster* expression atlas. Nucleic Acids Research 50, D1010–D1015 (2022).

14. Cao, J. et al. Insight into Insulin Secretion from Transcriptome and Genetic Analysis of Insulin-Producing Cells of Drosophila. Genetics 197, 175–192 (2014).

15. Söderberg, J. A. E., Carlsson, M. A. & Nässel, D. R. Insulin-Producing Cells in the Drosophila Brain also Express Satiety-Inducing Cholecystokinin-Like Peptide, Drosulfakinin. Front Endocrinol (Lausanne) 3, 109 (2012).

16. Park, S., Bustamante, E. L., Antonova, J., McLean, G. W. & Kim, S. K. Specification of Drosophila Corpora Cardiaca Neuroendocrine Cells from Mesoderm Is Regulated by Notch Signaling. PLoS Genet 7, e1002241 (2011).

17. Kim, S. K. & Rulifson, E. J. Conserved mechanisms of glucose sensing and regulation by Drosophila corpora cardiaca cells. Nature 431, 316–320 (2004).

18. Brogiolo, W. et al. An evolutionarily conserved function of the Drosophila insulin receptor and insulin-like peptides in growth control. Curr Biol 11, 213–221 (2001).

19. Cao, C. & Brown, M. R. Localization of an insulin-like peptide in brains of two flies. Cell Tissue Res 304, 317–321 (2001).

20. Rulifson, E. J., Kim, S. K. & Nusse, R. Ablation of insulin-producing neurons in flies: growth and diabetic phenotypes. Science 296, 1118–1120 (2002).

21. Corl, A. B., Rodan, A. R. & Heberlein, U. Insulin signaling in the nervous system regulates ethanol intoxication in Drosophila melanogaster. Nat Neurosci 8, 18–19 (2005).

22. Yamaguchi, S. T., Tomita, J. & Kume, K. The regulation of circadian rhythm by insulin signaling in Drosophila. 2022.04.25.489482 Preprint at 10.1101/2022.04.25.489482 (2022).

23. Bisen, R. S., Iqbal, F. M., Cascino-Milani, F., Bockemühl, T. & Ache, J. M. Nutritional state-dependent modulation of Insulin-Producing Cells in Drosophila. eLife 13, (2024).

24. Oyeyinka, A. et al. Corazonin Neurons Contribute to Dimorphic Ethanol Sedation Sensitivity in Drosophila melanogaster. Front. Neural Circuits 16, 702901 (2022).

25. Remy, N. Q., Guevarra, J. A. & Vonhoff, F. J. Food supplementation with wheat gluten leads to climbing performance decline in Drosophila melanogaster. MicroPubl Biol 2022, (2022).

26. Lee, P.-T. et al. A gene-specific T2A-GAL4 library for Drosophila. eLife 7, e35574 (2018).

27. Gabrawy, M. M. et al. Dual treatment with kynurenine pathway inhibitors and NAD^+^ precursors synergistically extends life span in *Drosophila*. Aging Cell 23, e14102 (2024).

28. Mishra-Gorur, K. et al. Mutations in KATNB1 cause complex cerebral malformations by disrupting asymmetrically dividing neural progenitors. Neuron 84, 1226–1239 (2014).

29. Veenstra, J. A., Agricola, H.-J. & Sellami, A. Regulatory peptides in fruit fly midgut. Cell Tissue Res 334, 499–516 (2008).

30. Itskov, P. M. et al. Automated monitoring and quantitative analysis of feeding behaviour in Drosophila. Nat Commun 5, 4560 (2014).

31. Ja, W. W. et al. Prandiology of Drosophila and the CAFE assay. Proceedings of the National Academy of Sciences 104, 8253–8256 (2007).

32. Gualtieri, C. et al. Dysregulation of Protein Kinase CaMKI Leads to Autism-Related Phenotypes in Synaptic Connectivity, Sleep, Sociality, and Aging-Dependent Degeneration in Drosophila. Biology (Basel) 14, 1228 (2025).

33. Vitaterna, M. H., Takahashi, J. S. & Turek, F. W. Overview of circadian rhythms. Alcohol Res Health 25, 85–93 (2001).

34. Ghosh, A. & Sheeba, V. VANESSA-Shiny Apps for Accelerated Time-series Analysis and Visualization of Drosophila Circadian Rhythm and Sleep Data. J Biol Rhythms 37, 222–231 (2022).

35. Hendricks, J. C. et al. Rest in Drosophila Is a Sleep-like State. Neuron 25, 129–138 (2000).

36. Shaw, P. J., Cirelli, C., Greenspan, R. J. & Tononi, G. Correlates of sleep and waking in Drosophila melanogaster. Science 287, 1834–1837 (2000).

37. Harbison, S. T., McCoy, L. J. & Mackay, T. F. C. Genome-wide association study of sleep in Drosophila melanogaster. BMC Genomics 14, 281 (2013).

38. Ponton, F., Chapuis, M.-P., Pernice, M., Sword, G. A. & Simpson, S. J. Evaluation of potential reference genes for reverse transcription-qPCR studies of physiological responses in Drosophila melanogaster. Journal of Insect Physiology 57, 840–850 (2011).

39. Broughton, S. J. et al. Longer lifespan, altered metabolism, and stress resistance in Drosophila from ablation of cells making insulin-like ligands. Proc Natl Acad Sci U S A 102, 3105–3110 (2005).

40. Devineni, A. V. et al. The genetic relationships between ethanol preference, acute ethanol sensitivity, and ethanol tolerance in *Drosophila melanogaster*. Fly 5, 191–199 (2011).

41. Devineni, A. V. & Heberlein, U. Acute ethanol responses in Drosophila are sexually dimorphic. Proc Natl Acad Sci U S A 109, 21087–21092 (2012).

42. Marini-Davis, A. et al. Juvenile hormone regulates the maturation of sexually dimorphic naive ethanol olfactory preference in Drosophila melanogaster. R Soc Open Sci 12, 242217 (2025).

43. Dakic, T. et al. The Expression of Insulin in the Central Nervous System: What Have We Learned So Far? IJMS 24, 6586 (2023).

44. Balleine, B. W., Liljeholm, M. & Ostlund, S. B. The integrative function of the basal ganglia in instrumental conditioning. Behav Brain Res 199, 43–52 (2009).

45. Cong, X. et al. Regulation of Sleep by Insulin-like Peptide System in Drosophila melanogaster. Sleep 38, 1075–1083 (2015).

46. Court, R. et al. Virtual Fly Brain—An interactive atlas of the Drosophila nervous system. Front. Physiol. 14, 1076533 (2023).

47. Catania, C., Binder, E. & Cota, D. mTORC1 signaling in energy balance and metabolic disease. Int J Obes 35, 751–761 (2011).

48. McBride, W. J. et al. Gene Expression within the Extended Amygdala of 5 Pairs of Rat Lines Selectively Bred for High or Low Ethanol Consumption. Alcohol 47, 10.1016/j.alcohol.2013.08.004 (2013).

49. Beckley, J. T. et al. The First Alcohol Drink Triggers mTORC1-Dependent Synaptic Plasticity in Nucleus Accumbens Dopamine D1 Receptor Neurons. J Neurosci 36, 701–713 (2016).

50. Nishimura, Y. et al. Genome-wide expression profiling of lymphoblastoid cell lines distinguishes different forms of autism and reveals shared pathways †. Human Molecular Genetics 16, 1682–1698 (2007).

51. Hodges, A. et al. Regional and cellular gene expression changes in human Huntington’s disease brain. Human Molecular Genetics 15, 965–977 (2006).

52. Brochier, C. et al. Quantitative gene expression profiling of mouse brain regions reveals differential transcripts conserved in human and affected in disease models. Physiological Genomics 33, 170–179 (2008).

53. Vonhoff, F. & Keshishian, H. Activity-Dependent Synaptic Refinement: New Insights from Drosophila. Front Syst Neurosci 11, 23 (2017).

54. Penney, J. et al. TOR Is Required for the Retrograde Regulation of Synaptic Homeostasis at the Drosophila Neuromuscular Junction. Neuron 74, 166–178 (2012).

55. Vonhoff, F. & Keshishian, H. In Vivo Calcium Signaling during Synaptic Refinement at the Drosophila Neuromuscular Junction. J Neurosci 37, 5511–5526 (2017).

56. Carrillo, R. A., Olsen, D. P., Yoon, K. S. & Keshishian, H. Presynaptic Activity and CaMKII Modulate Retrograde Semaphorin Signaling and Synaptic Refinement. Neuron 68, 32–44 (2010).

57. Vonhoff, F. & Keshishian, H. Cyclic nucleotide signaling is required during synaptic refinement at the Drosophila neuromuscular junction. Dev Neurobiol 77, 39–60 (2017).

58. Bai, Y. et al. Protein Kinase A Is a Master Regulator of Physiological and Pathological Cardiac Hypertrophy. Circulation Research 134, 393–410 (2024).

59. Mockett, B. G. et al. Calcium/Calmodulin-Dependent Protein Kinase II Mediates Group I Metabotropic Glutamate Receptor-Dependent Protein Synthesis and Long-Term Depression in Rat Hippocampus. J. Neurosci. 31, 7380–7391 (2011).

60. Polcowñuk, S., Yoshii, T. & Ceriani, M. F. Decapentaplegic Acutely Defines the Connectivity of Central Pacemaker Neurons in *Drosophila*. J. Neurosci. 41, 8338–8350 (2021).

61. Robles-Murguia, M. et al. Muscle-derived Dpp regulates feeding initiation via endocrine modulation of brain dopamine biosynthesis. Genes Dev. 34, 37–52 (2020).

62. Vo, H. D. L. & Lovely, C. B. Ethanol Induces Craniofacial Defects in Bmp Mutants Independent of nkx2.3 by Elevating Cranial Neural Crest Cell Apoptosis. Biomedicines 13, 755 (2025).

63. Takahama, K. et al. Pan-neuronal knockdown of the c-Jun N-terminal Kinase (JNK) results in a reduction in sleep and longevity in Drosophila. Biochem Biophys Res Commun 417, 807–811 (2012).

64. Kapfhamer, D. et al. JNK pathway activation is controlled by Tao/TAOK3 to modulate ethanol sensitivity. PLoS One 7, e50594 (2012).

65. Jee, C. & Batsaikhan, E. JNK Signaling Positively Regulates Acute Ethanol Tolerance in C. elegans. IJMS 25, 6398 (2024).

66. Berke, B., Wittnam, J., McNeill, E., Van Vactor, D. L. & Keshishian, H. Retrograde BMP signaling at the synapse: a permissive signal for synapse maturation and activity-dependent plasticity. J Neurosci 33, 17937–17950 (2013).

67. Berke, B., Le, L. & Keshishian, H. Target-dependent retrograde signaling mediates synaptic plasticity at the *Drosophila* neuromuscular junction. Developmental Neurobiology 79, 895–912 (2019).

68. Baines, R. A. Synaptic Strengthening Mediated by Bone Morphogenetic Protein-Dependent Retrograde Signaling in the *Drosophila* CNS. J. Neurosci. 24, 6904–6911 (2004).

69. McCabe, B. D. et al. The BMP Homolog Gbb Provides a Retrograde Signal that Regulates Synaptic Growth at the Drosophila Neuromuscular Junction. Neuron 39, 241–254 (2003).

70. Hartwig, C. L., Worrell, J., Levine, R. B., Ramaswami, M. & Sanyal, S. Normal dendrite growth in *Drosophila* motor neurons requires the AP-1 transcription factor. Developmental Neurobiology 68, 1225–1242 (2008).

71. Vonhoff, F., Kuehn, C., Blumenstock, S., Sanyal, S. & Duch, C. Temporal coherency between receptor expression, neural activity and AP-1-dependent transcription regulates *Drosophila* motoneuron dendrite development. Development 140, 606–616 (2013).

72. Freeman, A. A. H., Syed, S. & Sanyal, S. Modeling the genetic basis for human sleep disorders in Drosophila. Commun Integr Biol 6, e22733 (2013).

