## Supplemental figures for "Integral membrane protein, *anchor*, is expressed in the *Drosophila* insulin-producing cells and is a novel modulator of homeostatic behaviors, including sleep, feeding, and sedation"

***Supplemental materials***

***Supplemental Figure 1: Un-merged DILP2 and nuclear mCherry under anchor GAL4 driver.*** Representative images from single fly z-stack (slices 1-5 from first row of images to last) (n = 3). Immunohistochemistry using anti-DILP2 antibody (green, left) to image the insulin-producing cells (IPC)s in *Anchor GAL4 x nuclear mcherry* (red, middle) progeny. Merge (right) Each of the 14 DILP2-expressing IPCs indicated by arrowhead and numbered from 1-14.

***Supplemental Figure 2: Anchor RNAi1 and Anchor RNAi2 knockdown validation.*** *A) Male and B) Female verification panneuronal anchor knockdown, using RT-qPCR in whole-head samples. RT-qPCR samples run in group of 3-5 biological replicates in technical triplicate.*

***Supplemental Figure 3: anchor does not affect light response and perception.*** *A) Schematic of Phototaxis Assay. Made in Biorender. E) Data from phototaxis assay, showing how many flies moved towards light source (N = 18-35). One run (N) includes 5 flies).*
